# Compositional knockoff filter for high-dimensional regression analysis of microbiome data

**DOI:** 10.1101/851337

**Authors:** Arun Srinivasan, Lingzhou Xue, Xiang Zhan

## Abstract

A critical task in microbiome data analysis is to explore the association between a scalar response of interest and a large number of microbial taxa that are summarized as compositional data at different taxonomic levels. Motivated by fine-mapping of the microbiome, we propose a two-step compositional knockoff filter (CKF) to provide the effective finite-sample false discovery rate (FDR) control in high-dimensional linear log-contrast regression analysis of microbiome compositional data. In the first step, we employ the compositional screening procedure to remove insignificant microbial taxa while retaining the essential sum-to-zero constraint. In the second step, we extend the knockoff filter to identify the significant microbial taxa in the sparse regression model for compositional data. Thereby, a subset of the microbes is selected from the high-dimensional microbial taxa as related to the response using a pre-specified FDR threshold. We study the asymptotic properties of the proposed two-step procedure, including both sure screening and effective false discovery control. We demonstrate the finite-sample properties in simulation studies, which show the gain in the empirical power while controlling the nominal FDR. The potential usefulness of the proposed method is also illustrated with application to an inflammatory bowel disease dataset to identify microbial taxa that influence host gene expressions.

## 1. Introduction

The human microbiome refers to all the microbes that live in and on the human body with their collected genome, which has been linked to many human health and disease conditions (Cho and Blaser, 2012; Wang and Jia, 2016; Mitchell et al., 2017; Schneider et al., 2020). The advent of next-generation sequencing technologies enables studying the microbiome composition via direct sequencing of microbial DNA without the need for laborious isolation and cultivation, which largely boosts research interests in the human microbiome (Turnbaugh et al., 2007). Due to the varying amount of DNA yielding materials across different samples, the count of sequencing reads can vary significantly from sample to sample. As a result, it is a common practice to normalize the raw sequencing read counts to relative abundances making the microbial abundances comparable across samples (Weiss et al., 2017). Besides the compositional constraint, the increasing availability of massive human microbiome datasets, whose dimensionality is much larger than its sample size, also poses new challenges to statistical analysis (Li, 2015).

A central goal in microbiome analysis is fine-mapping of the microbiome to identify microbial taxa that are associated with a specific response of interest (e.g., body mass index or a host genomic/genetic feature). In general, existing methods of fine-mapping the microbiome fall into two main categories: marginal approach and joint approach. The marginal approach usually casts the fine-mapping problem into the microbiome-wide multiple testing framework by examining the marginal association between each microbial taxon and the response followed by multiple testing corrections to identify taxa with adjusted p-values below a certain FDR threshold as important ones that influence the response (Wang and Jia, 2016; Xiao, Chao, and Chen, 2017). The marginal approach is often limited to microbiome data analysis due to the following two reasons. First, it tends to have low detection power due to the heavy burden of multiple testing adjustment inherent from the high-dimensional nature of microbiome data (Li, 2015). Second, it fails to account for the simplex nature of compositional data and may suffer from spurious negative correlations imposed by the fact that relative abundances across all taxa must sum to one within a given microbiome sample. As a consequence, traditional FDR control procedures (Benjamini and Hochberg, 1995) may not work for microbiome-wide multiple testing (Hawinkel et al., 2017).

On the other hand, a joint microbiome selection approach usually models all taxa collectively using penalized regression (Chen and Li, 2013; Lin et al., 2014). These joint approaches achieve fine-mapping of the microbiome via variable selection, yet they have no guarantee on the false discoveries among the selected microbiome features. This is probably because the number of microbial features in the joint regression model is much larger than the sample size, and it is difficult to obtain a p-value measuring the significance of the association between the outcome and each microbial feature. Yet, a canonical FDR control approach in general needs to plug p-values into a certain multiple testing procedure (Benjamini and Hochberg, 1995). Without FDR control, existing joint microbiome fine-mapping methods can produce less reliable discoveries and would probably lead to costly and fruitless downstream validation and functional studies (Wang and Jia, 2016; Hawinkel et al., 2017).

To address the potential limitations in existing marginal and joint approaches, a new method in a joint regression framework to select microbial taxa with finite-sample FDR control is desired. In the statistics literature, the FDR control can be achieved via the knockoff filter framework, in which a dummy knockoff copy of the original design matrix with the same covariance structure has been constructed and flagged as false positives to facilitate FDR-controlled variable selection (Barber and Candès, 2015). However, it has been observed, in the literature of many other statistical inference methods (e.g., regression-based modeling, two-sample testing, and statistical causal mediation analysis), that applying classic statistical methods to analyze microbiome composition data is usually underpowered and sometimes can render inappropriate results (Aitchison, 2003; Shi, Zhang, and Li, 2016; Cao, Lin, and Li, 2017; Sohn and Li, 2019; Lu, Shi and Li, 2019; Zhang et al., 2019). Thus, new FDR-controlled variable selection methods are desired for microbiome compositional data.

Following the pioneering work of Aitchison and Bacon-shone (1984), we model all taxa jointly in a linear log-contrast model to address the compositional nature of data and propose a two-step regression-based FDR-controlled variable selection procedure named compositional knockoff filter (CKF) to identify response-associated taxa. In the first step, we introduce the compositional screening procedure (CSP) as a new method of variable screening for high-dimensional microbiome data subject to the compositional constraint. In the second step, we apply the fixed-X knockoff procedure (Barber and Candès, 2015) to the reduced model in the first screening step. The theoretical properties of the novel compositional screening procedure are investigated. Using numerical studies, we demonstrate that the proposed method can jointly assess the significance of microbial covariates while also theoretically ensuring finite-sample FDR control. The proposed method will greatly benefit downstream microbiome functional studies by enhancing the reproducibility and reliability of discovery results in microbiome association studies.

Our primary contributions are summarized as follows. First, we introduce the CSP to screen true signals from high-dimensional compositional data and theoretically verify that CSP attains the desirable sure screening property under mild assumptions. As demonstrated in thorough simulation, the newly proposed CSP yields a much higher likelihood of attaining all true signals compared to some existing methods that do not account for the compositional nature. Second, by leveraging the high-dimensional knockoff filter framework (Barber and Candès, 2019), we avoid the non-trivial sequential conditional independent pairs algorithm of model-X knockoffs (Candès et al., 2018) and provide an alternative CKF approach to ensure strong finite-sample FDR control for microbial taxa selection. Construction of model-X knockoff features through methods such as the sequential conditional independent pairs algorithm (Candès et al., 2018) requires both complete knowledge of the joint distribution of the microbiome design matrix and repeated derivation of the conditional distributions, that are non-trivial for many non-Gaussian distributions such as Dirichlet-multinomial and logistic normal, which are frequently used in modeling microbiome data (Aitchison, 2003; Chen and Li, 2013; Lin et al., 2014; Tang and Chen, 2018; Harrison et al., 2020). While the development of methods to construct exact or approximate knockoff features for a broader class of distributions is a promising area of active research (Bates et al., 2019; Romano, Sesia, and Candès, 2019), the robustness of how model-X knockoff to the departure of the joint distribution from multivariate Gaussian is currently unknown. To this end, the proposed CKF with finite-sample FDR control guarantee is appealing through versatility for microbiome taxa selection.

The rest of this paper is organized as follows. We propose the methodology of compositional knockoff filter in Section 2. The theoretical properties of the compositional screening procedure are investigated in Section 3. The numerical properties are demonstrated through simulation studies in Section 4 and application to a microbiome data set collected from an inflammatory bowel disease study in Sections 5. Technical proofs and additional numerical evaluations are deferred to the online supporting information.

## 2. Compositional Knockoff Filter

This section presents the compositional knockoff filter to perform FDR-controlled variable selection analysis for microbiome compositional data. The proposed method aims to address the high-dimensional compositional nature of microbiome data (i.e., *p > n*). To this end, we follow the philosophy of recycled fixed-X knockoff procedure (Barber and Candès, 2019) to develop a new two-step procedure for high-dimensional compositional data, which consists of a compositional screening step and then a subsequent selection step. After introducing the log-contrast model in Section 2.1, we will present the screening step in Section 2.2 and the selection step in Section 2.3.

### 2.1 Log-Contrast Model

We use the log-contrast model (Aitchison and Bacon-shone, 1984) for joint microbiome regression analysis. Let **Y** *∈* ℝ^*n*^ denote the response vector and **X** *∈* ℝ^*n×p*^ denote a matrix of microbiome compositions. By structure of the microbiome compositional components, each row of **X** must individually sum to 1. Thus **X** is not of full rank, leading to identifiability issues for the regression parameters. In order to account for this structure, the log-linear contrast model is often used for compositional data (Aitchison, 2003; Lin et al., 2014). We assume that *X*_*ij*_ *>* 0 by replacing the zero proportions by a tiny pseudo positive value as routinely performed in practice (Lin et al., 2014; Shi et al., 2016; Cao et al., 2017; Lu et al., 2019; Zhang et al., 2019). Let **Z**^*p*^ *∈* ℝ^*n×*(*p−*1)^ be the log-ratio transformation of **X**, where 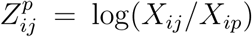 and *p* denotes the reference covariate. The linear log-contrast model is formulated as **Y** = **Z**^*p*^***β***_*\p*_ + *ε*, where ***β***_*\p*_ is the vector of (*p −* 1) coefficients (*β*_1_, *β*_2_, *…, β*_*p−*1_) and error *ε ∼ N* (0, *σ*^2^**I**). To avoid picking a reference component for better model interpretability, the log-contrast model is often reformulated into a symmetric form with a sum-to-zero constraint (Lin et al., 2014). That is,

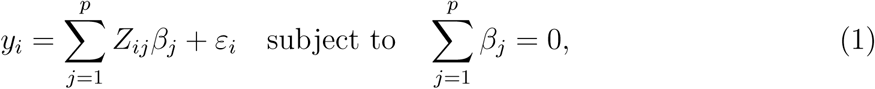

where **Z** *≡ {Z*_*ij*_*}* is the *n × p* log-composition matrix with *Z*_*ij*_ = log(*X*_*ij*_) and ***β*** *≡* (*β*_1_, *β*_2_, *…, β*_*p*_)^*′*^ are the regression coefficients for microbiome covariates. For ease of presentation, model (1) does not explicitly include other covariates, but all the results in the rest of this article still hold in presence of other covariates.

### 2.2 Compositional Screening Procedure

As the fixed-X knockoff requires that *n ⩾* 2*p*, screening the predictor set to a low-dimensional setting is necessary for the analysis of high-dimensional compositional data. Let *n*_0_ denote the number of samples to use for screening and *n*_1_ denote the remaining observations, where *n* = *n*_0_ + *n*_1_. We randomly split the original data (**Z, Y**) into (**Z**^(0)^, **Y**^(0)^) and (**Z**^(1)^, **Y**^(1)^), where 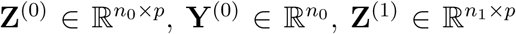 and 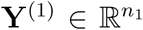. By ensuring that **Z**^(0)^ and **Z**^(1)^ are disjoint, we are able to implement a recycling step to reuse the original screening data **Z**^(0)^, in order to increase the selection power. To this end, we first use the sub-data (**Z**^(0)^, **Y**^(0)^) to perform the screening and obtain a subset of features Ŝ_0_ ⊂ {1, *…, p*} such that 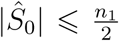, where |Ŝ_0_| denotes the cardinality of set Ŝ_0_. Throughout this paper, we always assume 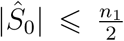 to ensure that we are able to construct the fixed-X knockoffs (Barber and Candès, 2015) for data (**Z**^(1)^, **Y**^(1)^) in the subsequent selection step. As the selection step further reduces the feature set after screening, we must ensure that true signals are not lost before the selection step. For this reason, we desire screening methods that attain the sure screening property (Fan and Lv, 2008). That is, with high probability, we desire the selection set estimated by the screening method of choice to contain all relevant features. It is popular to perform screening using Pearson correlation (Fan and Lv, 2008; Fan and Song, 2010; Xue and Zou, 2011) or distance correlation (Li, Zhong and Zhu, 2012). Despite that both marginal correlations-based screening methods enjoy the sure screening property asymptotically, these methods do not account for the compositional nature of microbiome data, which might lead to inefficient inference. We will further demonstrate this issue in the simulation studies of Section 4.1.

To account for the compositional structure, we introduce the novel compositional screening procedure to improve the efficiency for screening microbiome compositional covariates. In general, best-subset selection is often used to identify the optimal *k* best features (Beale, Kendall and Mann, 1967). In our log-contrast model, the best-subset selection problem can be expressed as a constrained sparse least-squares estimation problem as follows:

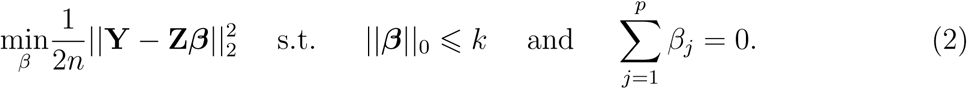

The proposed compositional screening problem (2) can also be viewed as maximizing the log-likelihood *ℓ*_*n*_(***β***) under the sparsity constraint ||***β***||_0_ ⩽ *k* (Xu and Chen, 2014). The choice of *k* is a fundamental question in many high-dimensional screening procedures. Practical domain knowledge may provide information on how sparse one believes the underlying signal is. Common choices for screening set size are often 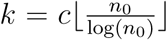 for some *c >* 0 (Fan and Lv, 2008; Li et al., 2012). However, as noted by Li et al. (2012), the choice of screening set size can be viewed as a tuning parameter within the model and concrete means to determine the screening set size are an area of future development. Although (2) is a NP-hard problem, the mixed integer optimization (MIO) allows us to approximately solve the global solution of the nonconvex optimization problem (2) in an efficient manner (Konno and Yamamoto, 2009; Bertsimas, King and Mazumder, 2016). Finally, we demonstrate in the Section 3 that the computed solution of (2) by MIO attains the desirable sure screening guarantees.

After screening, the model reduces to 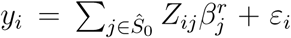 subject to 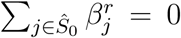. Comparing it to the original log-contrast model (1), the regression coefficients in the reduced model 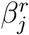 does not necessarily match *β*_*j*_ in the original model. To solve this discrepancy, we propose a normalization procedure 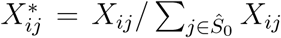 for *j ∈ Ŝ*_0_ and for an abuse of notation, we still use 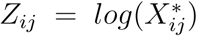 to denote the design matrix to be used in the subsequent selection step. Details about this normalization is available at Section S.1 of the online supporting information.

### 2.3 Controlled Variable Selection

Let 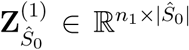 denote the columns of **Z**^(1)^ corresponding to *Ŝ*_0_, the selected set from the *computed* solution of (2), and we delineate this from the selection set from the *global* solution of (2) which we instead denote as *Ŝ*_0_. The knockoff matrix 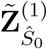 is constructed using 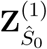 following the fixed-X knockoff framework (Barber and Candès, 2015). The primary assumption of the fixed-X framework is that the burden of knowledge is placed on the design and **Z**^(1)^ is assumed to be fixed and the response is generated via a linear Gaussian model. Notably, the fixed-X knockoff places no assumptions on knowing the noise level. Thus, a key appeal of the knockoff filter is the relative lack of strong assumptions needed for theoretical finite-sample control to hold. We refer to Barber and Candès (2015) for a review of the construction of knockoff matrix and for a deeper study into the assumptions needed by the knockoff filter. The use of the screening step allows us to apply the fixed-X knockoff framework in the high-dimensional setting. While fixed-X knockoffs traditionally require a low-dimensional regime, the screening step first reduces the effective dimension to one of size at most 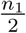. Thus 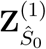 is of dimension at most 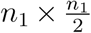. As the knockoff matrix is constructed on 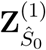 alone, this satisfies the dimensionality requirements for the construction of the fixed-X knockoff matrix 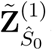. To further boost the power of the procedure, we followed data recycling mechanism outlined in Barber and Candès (2019), we construct the recycled knockoff matrix as

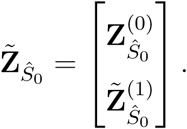

Note that we treat (**Z**^(0)^, **Y**^(0)^) as fixed after the screening step and the first part of knockoff copies are exact copies under the recycling scheme (Barber and Candès, 2019).

We now run the knockoff regression procedure using 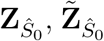, and **Y**. In particular, we first append the screened original and knockoff matrices to create an augmented design matrix 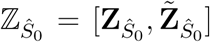. This augmented design matrix is of dimension 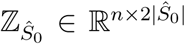 where the first |*Ŝ*_0_| features are the original covariates and the remaining |*Ŝ*_0_| features are the knockoff copies. With this new augmented design matrix, we solve the following Lasso problem:

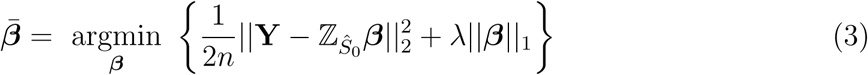

where 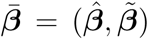 is a vector appending the coefficients of original features and knockoff features. Other penalties such as the folded concave penalties (Fan and Li, 2001; Fan et al., 2014) may also be used for the purpose of variable selection. For ease of presentation, we focus on the Lasso problem (3), as existing methods (Lin et al., 2014) do not provide a rigorous FDR control on the selected variables. Comparing to previous problems (1) and (2), we no longer require a sum-to-zero constraint in our augmented Lasso problem (3). This is because, by adding |*Ŝ*_0_| knockoff features in the augmented design matrix 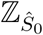, the corresponding microbiome data matrix 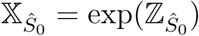 is no longer compositional in nature.

The above optimization problem (3) is performed over the entire Lasso path and provides a set of Lasso coefficients denoted by 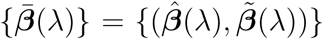. Based on 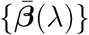, we next calculate the knockoff statistic *W*_*j*_, which measures evidence against the null hypothesis *β*_*j*_ = 0 for each *j ∈ Ŝ*_0_. For the scope of this paper we use the Lasso signed lambda max statistic (LSM). Let 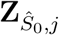 denote original covariate *j* and 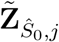 denote knockoff covariate *j*:

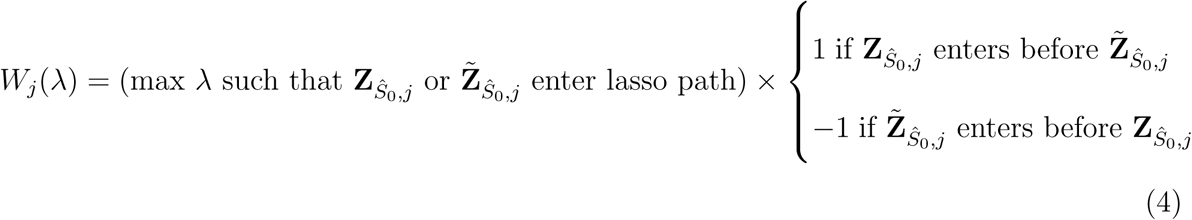

A large and positive *W*_*j*_ would suggest strong evidence that the original feature is significantly outcome-associated as an important feature tends to remain longer in lasso path as *λ* increases. Similarly, a negative or zero *W*_*j*_ value would indicate that the covariate tends to be noise. Thus, *W*_*j*_ is used to calculate the data-dependent knockoff thresholds that ensure finite sample FDR-controlled variable selection. The choice of an *ℓ*_1_-penalization problem used in the selection step is driven by the need to accurately compute a solution path for construction of the knockoff statistic. As a comparison, the solution of *ℓ*_0_-penalization problem (2) via MIO does not yield a solution path and is unsuitable for knockoff test statistic construction. Finally, both the standard knockoff and knockoff+ thresholds are considered for the purpose of selection:

Knockoff Threshold:

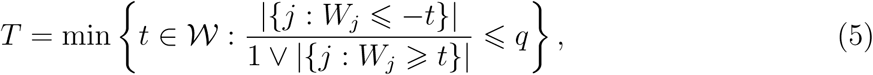

Knockoff+ Threshold:

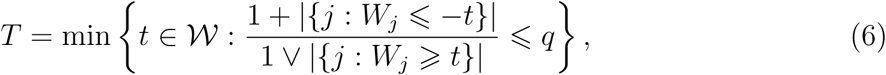

where *q ∈* [0, 1] is the user-specified nominal FDR level, 𝒲 = *{*|*W*_*j*_| : *j ∈ Ŝ*_0_*}\{*0} are the unique non-zero values of |*W*_*j*_|’s (*T* = +*∞* if 𝒲 is empty) and *a* ∨ *b* denotes the maximum of *a* and *b*. Once this threshold has been calculated, we select covariates *Ŝ* = *{j* : *W*_*j*_ *⩾ T*}. Depending on the threshold being used, we term this FDR-control variable selection procedure as either compositional knockoff filter (CKF) or compositional knockoff filter+ (CKF+). For completeness, we summarize the proposed CKF procedures in Algorithm 1.

## 3. Theoretical Properties

In this section, we first present the theoretical properties of the proposed compositional screening procedure and show that the computed solution from solving the constrained sparse maximum likelihood problem (2) via the mixed integer optimization attains the desired sure screening property. We then summarize the theoretical properties of the proposed compositional knockoff filter method. Leveraging the framework of high-dimensional knockoff filter (Barber and Candès, 2019), we verify that CKF/CKF+ attain finite sample FDR control under the compositional constraint. The main results are presented in this main text and details on the proof to establish these theoretical properties is available through Section S.3 of the online supporting information.

### Algorithm 1 Compositional Knockoff Filter (CKF)

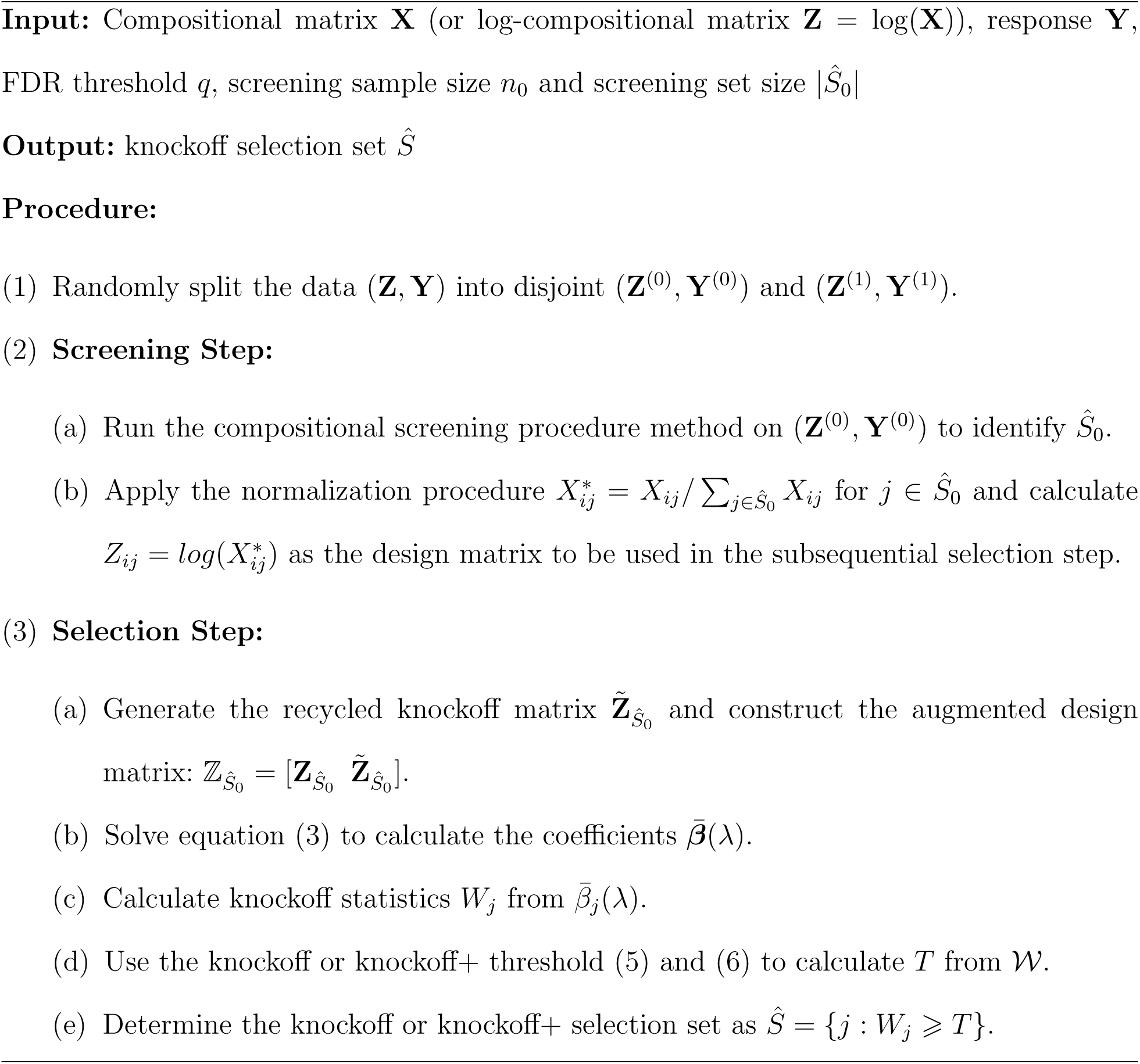

### 3.1 Theoretical Properties of Compositional Screening

We will show in this section that the compositional screening procedure attains the sure screening property. For ease of presentation, some notation is introduced first. Let *s* denote an arbitrary subset of *{*1, *…, p*} corresponding to a sub-model with coefficients *β*_*s*_, and *S*^*\**^ be the true model with *p*^*\**^ nonzero coefficients, with corresponding true coefficient vector ***β***^*\**^. Let *Ŝ*_0_ denote the computed screened sub-model after applying the compositional screening procedure. Assume that *Ŝ*_0_ retains at most *k* features with 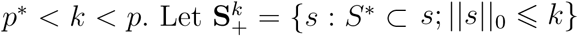 denote the set of all overfit models and 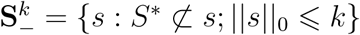 denote the set of underfit models. We will show that the compositional screening procedure does not miss true signals with high probability. That is:

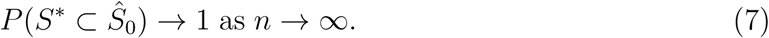

For the technical aspects of our sure-screening proof to hold, we make the following assumptions (1-4), encompassing requirements on the dimension, signal strength and microbiome design matrix:

Assumption 1: log(*p*) = *O*(*n*^*m*^) for some 0 ⩽ *m <* 1.

Assumption 2: There exists *w*_1_ *>* 0 and *w*_2_ *>* 0 and non-negative constants *τ*_1_ and *τ*_2_ such that 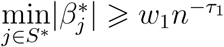 and 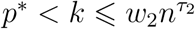.

Assumption 3: There exist constants *c*_1_ *>* 0 and *δ*_1_ *>* 0 such that for sufficiently large *n* such that 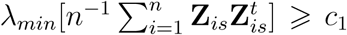 for 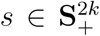 and 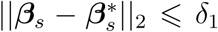, where *λ*_*min*_[*M*] denotes the smallest eigenvalue of the matrix *M*, and **Z**_*is*_ = (*Z*_*ij*_)_*j∈s*_.

Assumption 4: There exist constants *c*_2_ *>* 0 and *c*_3_ *>* 0 such that |*Z*_*ij*_| *⩽ c*_2_ and 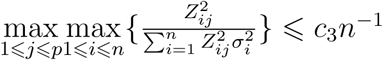 when *n* is sufficiently large, where 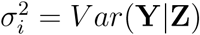.

Assumption 1 places a weak restriction on *p* and *n* of the data, which is very likely to be met in many microbiome studies (Wang and Jia, 2016). Assumption 2 places a restraint on the minimum strength of true signals, such that they are discoverable. This assumption is common for statistical screening and variable selection methods (Fan and Lv, 2008; Fan and Song, 2010; Lin et al., 2014). Assumption 3 corresponds to the UUP condition (Candes and Tao, 2007) which controls the pairwise correlations between the columns of **Z**. This condition is prevalent across many high-dimensional variable selection methods such as the Dantzig selector (Candes and Tao, 2007), SIS-DS (Fan and Lv, 2008), forward regression (Wang, 2009), and the sparse MLE (Xu and Chen, 2014). We have conducted numerical studies to evaluate the applicability of Assumption 3 in the context of microbiome data settings in Section S.2 of the online supporting information. Finally, as noted by Xu and Chen (2014) and Chen and Chen (2012), Assumption 4 likely will hold for a wide class of design matrices as long 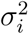 is not degenerate. In Section S.2 of the online supporting information, we illustrate the validity of Assumption 4 on the mucosal microbiome data analyzed later in this paper. Further details on these assumptions are available in Section S.2 of the online supporting information. Under Assumptions 1–4, Theorem 1 shows that the proposed compositional screening procedure attains the sure screening property. The proof of Theorem 1 relies on two key lemmas which are presented first.

#### Lemma 1

*Let* 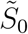 *denote the index set of screened features from the global solution of the constrained sparse maximum-likelihood estimation problem* (2), *where* 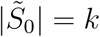. *Let* 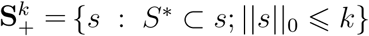. *Assume that Assumptions 1-4 hold and* 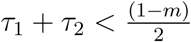. *Then:*

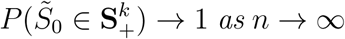

Lemma 1 ensures that the model selected by the solution of the constrained sparse maximum-likelihood estimation will be in the set of overfit models with high-probability. Thus, this ensures no signals are lost during screening. In other words, the global solution of the constrained sparse maximum-likelihood estimation problem attains the sure screening property.

#### Lemma 2

*Let* 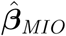 *denote the computed coefficient magnitudes of the model selected by the compositional screening procedure through mixed integer optimization and* 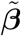 *denote the coefficients of the global solution of the constrained sparse maximum likelihood problem. Given ε >* 0, *then:*

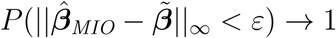

Lemma 2 demonstrates that the computed solution of the compositional screening procedure through mixed integer optimization converges to the global solution of the constrained sparse maximum likelihood problem with high probability. By combining Lemma 1 and Lemma 2, it follows that the computed solution attains the sure screening property. This result is presented in Theorem 1.

#### Theorem 1

*Given we have n independent observations with p possible features. Assume that Assumptions 1-4 hold and* 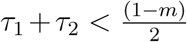. *Let Ŝ*_0_ *denote the computed screened set from the compositional screening procedure where p*\* *<* |*Ŝ*_0_| *< p. Then:*

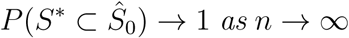

Theorem 1 allows us to claim that the compositional screening procedure will not lose any signals during screening with high probability. In summary, the compositional screening procedure accounts for the compositional constraint and also ensures the screening power.

### 3.2 FDR Control Properties of Compositional Knockoff Filter

In this section, we briefly outline the FDR control properties of CKF. In order to control FDR, the knockoff statistic must obey the *anti-symmetry* and *sufficiency* properties while the design matrix and response must satisfy both the *Pairwise Exchangeability for the Response Lemma* and *Pairwise Exchangeability for the Features Lemma* (Barber and Candès, 2015). In this paper we primarily focus on the LSM knockoff statistic (4) which has been shown to satisfy the *anti-symmetry* and *sufficiency* properties (Barber and Candès, 2019). Therefore, the FDR control properties of CKF are a direct consequence of the FDR control theory outlined in Barber and Candès (2015) and Barber and Candès (2019) and we reiterate the FDR control property here for posterity. As we have validated that the CSP attains the sure-screening property, the compositional knockoff+ threshold ensures finite sample FDR control as stated in the following Theorem 2.

#### Theorem 2

*For q ∈* [0, 1], *the knockoff+ method with data-recycling ensures:*

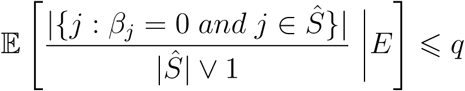

*where S denotes the index set of selected coefficients through the compositional knockoff+ procedure, E denotes the event {S*^*\**^ *⊂ Ŝ*_0_*}. The expectation is over the Gaussian noise vector ε and* **Z** *and* 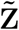 *are fixed.*

Theorem 2 demonstrates that CKF+ controls the FDR at a user-specified level *q*, after conditioning on the results of the screening procedure. By the argument in Theorem 2 of Barber and Candès (2019), if a proper screening procedure which attains the sure screening property (such as our proposed compositional screening procedure through mixed integer optimization) is implemented in the screening step, FDR is controlled even without conditioning on *E*.

#### Remark 1

Given the above exchangeability results and the previous theorems, the standard knockoff threshold controls a modified form of false discovery rate (Barber and Candès, 2015). In particular, for *q ∈* [0,1], the knockoff method en ures:

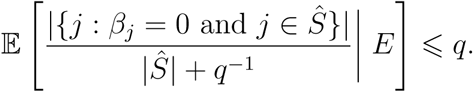

Compared with the formula in Theorem 2, the additional *q*^*−*1^ in the denominator sometimes favors a larger selected set *Ŝ* in CKF compared to CKF+. But when the selected set *Ŝ* is relatively large or when the nominal FDR threshold *q* is relatively large, the difference between CKF and CKF+ vanishes as *q*^*−*1^ has little effect compared to |*Ŝ*| under such scenarios.

## 4. Simulation Studies

We conducted two sets of simulation studies (screening simulation and selection simulation) to evaluate numerical performance of the proposed CKF methods. In the screening simulation, we evaluated the sure screening property of the proposed CSP. We compared CSP to two other popular statistical screening procedures in literature: one based on Pearson correlation/PC (Fan and Lv, 2008) and the other based on distance correlation/DC (Li et al., 2012). We considered a sample size of *n*_0_ = 100 and screening set size 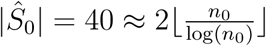. In the selection simulation, we evaluated the selection performance of CKF methods. For comparison, we also consider other methods that are widely used for microbial taxa selection. One is the compositional Lasso (Lin et al., 2014) and the other is the marginal method which examines one taxon at a time followed by the Benjamini-Hochberg procedure for FDR control (Benjamini and Hochberg, 1995; Paulson et al., 2013; Parks et al., 2014). We also compared the proposed CKF to the original model-X knockoff filter method (Candès et al., 2018) in this selection simulation. To mimic a real dataset analyzed later in this paper, we considered sample size *n* = 250 and number of microbiome covariates *p* = 400 in the selection simulations. Among these *n* = 250 samples, a randomly selected sub-sample with *n*_0_ = 100 observations were used in the first screening step and the rest *n*_1_ = 150 observations were used for the selection step.

Two schemes have been used to generate the microbiome compositional design matrix used in both simulations. The first scheme was to generate microbiome counts from the Dirichlet-multinomial (DM) distribution, whose parameters were estimated from a real microbiome data set following previous designs (Zhao et al., 2015). The library size of each sample was randomly simulated from a negative binomial distribution with a mean parameter of 10000 and dispersion parameter of 25. Raw zero counts were first replaced by a pseudo count of 0.5, as commonly suggested in microbiome data analysis (Lin et al., 2014; Cao et al., 2017; Weiss et al., 2017; Lu et al., 2019; Zhang et al., 2019) and then counts were transformed to relative abundances. The second scheme for generating microbiome compositional data was to use the logistic normal (LN) distribution, which is also widely used to generate compositional data (Aitchison, 2003; Lin et al., 2014; Cao et al., 2017). Following a previous design (Lin et al., 2014), we first simulated an intermediate *n × p* data matrix **M** from multivariate normal distribution *N*_*p*_(***µ*, Σ**), where *µ*_*i*_ = 1 and Σ_*ij*_ = 0.5^|*i−j*|^ for *i, j* = 1, *…, p*. Then, we calculated the log-composition design matrix as 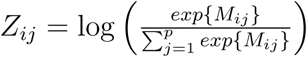 for *i* = 1, *…, n, j* = 1, *…, p*.

Next, we varied the sparsity levels |*S*^*\**^| ∈ {15, 20, 25, 30} and set the first 30 entries ***β***_1:30_ of the whole regression coefficient vector ***β***_1:400_ as: ***β***_1:30_ = (*−*3, 3, 2.5, *−*1, *−*1.5; 3, 3, *−*2, *−*2, *−*2; 1, *−*1, 3, *−*2, *−*1; *−*1, 1, 2, *−*1, *−*1; 3, 3, *−*3, *−*2, *−*1; 3, 3, *−*3, *−*2, *−*1). The remaining regression coefficients ***β***_31:400_ were all set to be zeros. We constructed the regression coefficients in the aforementioned way such that 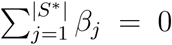, for each |*S*^*\**^| ∈ {15, 20, 25, 30}. Under this scheme, it is easy to check that the coefficient vector always satisfies the sum-to-zero constraint under each of the four sparsity levels. Finally, we simulated the response vector **Y** from 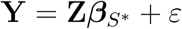, where 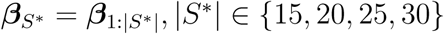 and *ε ∼ 𝒩* (0, *I*).

### 4.1 Screening Simulation

We first applied the three screening methods (CSP, PC, DC) to the simulated data to evaluate the screening accuracy by calculating the proportion of true features being selected in the screened set, |*Ŝ*_0_ *∩ S*^*\**^|*/*|*S*^*\**^|, where *Ŝ*_0_ is the screening set and *S*^*\**^ is the set of covariates with true non-zero coefficients in the log-contrast model. The results on screening accuracy of different methods are summarized in Table 1.

**Table 1.**
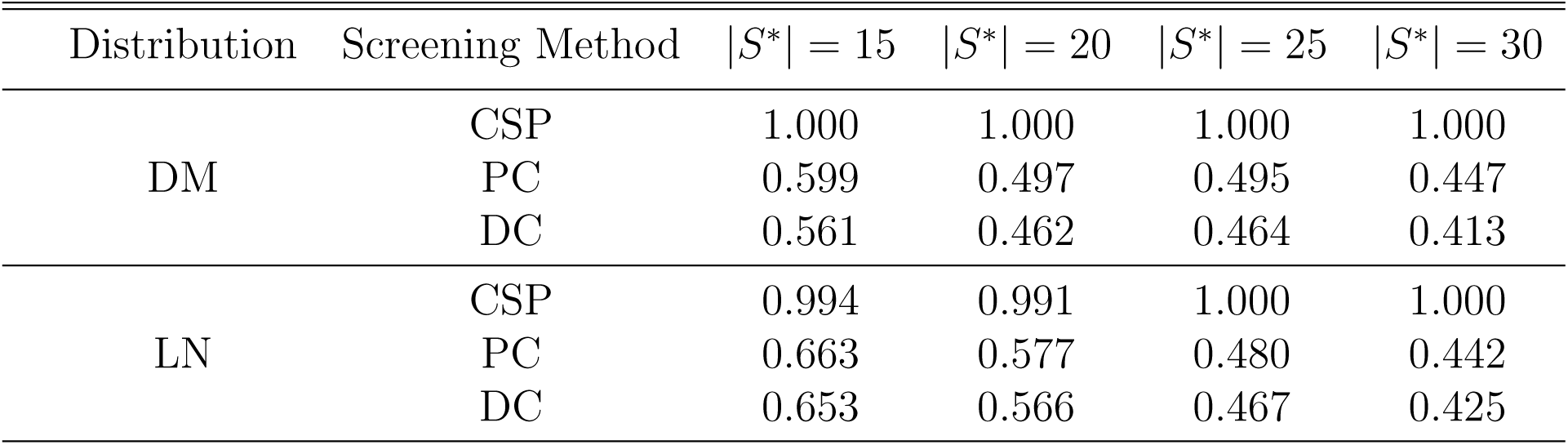
Average screening proportions of true signals based on 200 replicates under the Dirichlet-multnomial (DM) distribution and logistic normal (LN) distribution.

| Distribution | Screening Method | $\vert S^*\vert = 15$ | $\vert S^*\vert = 20$ | $\vert S^*\vert = 25$ | $\vert S^*\vert = 30$ |
| --- | --- | --- | --- | --- | --- |
| DM | CSP | 1.000 | 1.000 | 1.000 | 1.000 |
|  | PC | 0.599 | 0.497 | 0.495 | 0.447 |
|  | DC | 0.561 | 0.462 | 0.464 | 0.413 |
| LN | CSP | 0.994 | 0.991 | 1.000 | 1.000 |
|  | PC | 0.663 | 0.577 | 0.480 | 0.442 |
|  | DC | 0.653 | 0.566 | 0.467 | 0.425 |

The proposed CSP has much better performance than the other two competing methods PC and DC (Fan and Lv, 2008; Li et al., 2012), which have been widely used in the statistical literature. This is another example that classic statistical methods may be inefficient for microbiome data without accounting for the compositional nature (Lin et al., 2014; Shi et al., 2016; Cao et al., 2017; Lu et al., 2019; Zhang et al., 2019). By incorporating the compositional constraint, the proposed CSP achieves the sure screening property for microbiome data as the proportion of true features retained in the screened set is always one based on Table 1. It is of note that the performance of screening is crucial to the subsequent selection inference. To show this, we have further conducted additional numerical studies to compare the performance of CKF and CKF+ with three different screening procedures at a target nominal FDR of 0.1. The results are reported in Table S4 and Table S5 in Section S.4 of the online Supporting Information.

### 4.2 Selection Simulation

In this section, we compared CKF/CKF+ to some existing methods including the model-X knockoff filter (KF) methods (Candès et al., 2018), compositional lasso (CL) method (Lin et al., 2014) and Benjamini-Hochberg (BH) procedure. The KF method places the burden of knowledge on knowing the complete conditional distribution of **Z**, and there is no algorithm that can generate model-X knockoffs for general distributions efficiently (Bates et al., 2019). Therefore, we employ the use of Gaussian model-X knockoffs as used previously (Candès et al., 2018) in this simulation. For the CL method, the optimal *λ* used in the compositional Lasso was determined through 10-fold cross-validation. As the number of microbial features is typically larger than the sample size in microbiome association studies, it is difficult to obtain joint association p-values for each microbial feature. We examined the association between the outcome and each microbial feature marginally and applied the Benjamini-Hochberg (BH) procedure to these marginal p-values to identify features significant under FDR of 0.1. To measure performance of different selection methods, empirical FDR and empirical power were calculated.

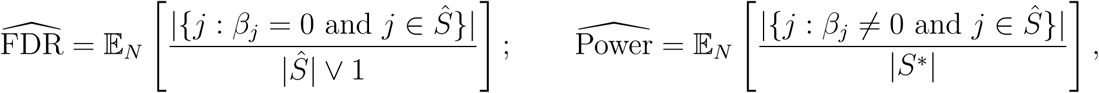

where 𝔼_*N*_ denotes the empirical average over *N* = 200 replicates. The results of empirical FDR and empirical power are reported in Table 2.

**Table 2.**
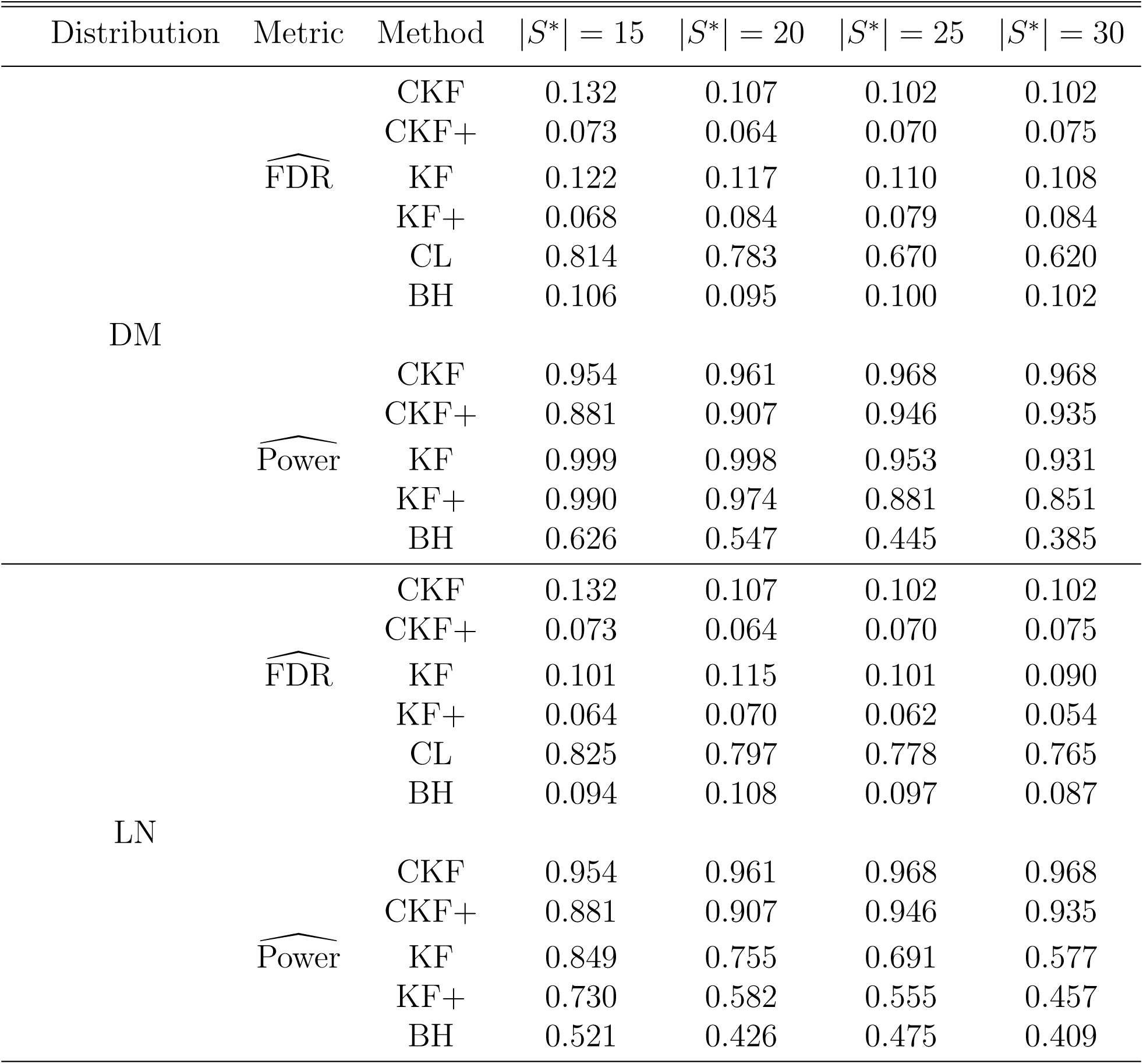
Empirical FDR and power under nominal FDR of 0.1 based on 200 replicates.

As observed from Table 2, CKF+, KF+ and BH can control the nominal FDR level, which is desired. CKF and KF yield slightly inflated FDR levels above the nominal rate, but this is expected as both KF and CKF are only guaranteed to control a modified version of the FDR (Remark 1 of Theorem 2). Finally, CL has a high empirical false discovery rate across all scenarios. The Lasso method has proven to be a versatile tool with appealing estimation and selection properties in the asymptotic setting (Tibshirani, 1996; Lin et al., 2014). Yet, its performance under finite sample setting is not guaranteed. Our results on CL is consistent with the fact that a relatively large number of false positives are reported in Table 1 of Lin et al. (2014). Despite being able to guarantee model selection consistency, CL tends to select more unnecessary variables to recover the true model.

Since CL has an extremely inflated FDR, it is not meaningful to compare its power to the other methods that can control FDR and hence power of CL is not reported. Comparing the empirical power of methods with FDR contrrol in Table 2, both CKF+ and KF+ are much more powerful than BH. This power gap is likely due to the fact that CKF+ and KF+ analyze the microbial covariates jointly, and the effectiveness of the marginal BH method deteriorates when the dimension (or multiple correction burden) is relatively high. Under DM distribution, KF+ achieves as higher power than CKF+ in sparse setting (|*S*^*\**^| = 15 or 20). However, CKF+ becomes more powerful over KF+ as the signal becomes denser (|*S*^*\**^| = 25 or 30). On the other hand, under LN distribution, the effectiveness of KF+ quickly deteriorates and CKF+ is much more powerful than KF+ based on Table 2. The KF+ method generated knockoffs based on an underlying Gaussian assumption on the covariates, and therefore its performance under the microbiome setting (e.g., Dirichlet-multinomial and logistic normal distributions considered in our simulations) may not be guaranteed. As a comparison, the proposed CKF+ method avoids the assumption on the joint distribution of design matrix and therefore is more robust to potential model misspecification. We limited the aforementioned discussions to CKF+ and KF+, while the same conclusions also apply when comparing CKF and KF.

Finally, we note that theoretically, the CKF method is only guaranteed to control a modified version of the FDR and usually has a higher FDR level than CKF+. In exploratory settings where FDR control is not at a premium, we suggest using CKF as the default for maximal power across all sparsity levels. In non-exploratory settings where users wish to have rigorous FDR control, we suggest the use of CKF+ as the default since CKF+ ensures theoretical finite sample FDR control and still attains high power in a majority of settings. To summarize, the proposed CSP enjoys the sure screening property, which is crucial to guarantee a high power of the downstream selection analysis. Our CKF methods successfully control the FDR of selecting outcome-associated microbial features in a regression-based manner which jointly analyzes all microbial covariates, while having the highest power detecting outcome-associated microbes. CKF methods are more robust than KF methods modelling the microbiome compositional data and are more powerful than the corresponding KF methods under most scenarios. Compared to CKF methods, Other existing methods may either be underpowered (BH) or render inappropriate results (CL) by having an inflated FDR than the nominal threshold.

## 5. Real Data Example

To further demonstrate the usefulness of our method, we apply it to a real data set obtained from a study examining the association between host gene expression and mucosal microbiome using samples collected from patients with inflammatory bowel disease (Morgan et al., 2015). The abundances of 7000 OTUs from *n* = 255 samples were measured using 16S rRNA gene sequencing and most to these 7000 species-level OTUs were in extremely low abundances with a large proportion of OTUs being simply singletons. As suggested in literature (Li, 2015), we aggregated these OTUs to genus and perform the analysis in the genus level, which may be more robust to potential sequencing errors. These 7000 OTUs belonged to *p* = 303 distinct genera, whose abundances were the microbial covariates of interest in our analysis.

It has been previously found that microbially-associated host transcript pattern is enriched for complement cascade genes, such as genes CFI, C2, and CFB (Morgan et al., 2015). Moreover, principal component-based enrichment analysis shows that host gene expression is inversely correlated with taxa *Sutterella, Akkermansia, Bifidobacteria, Roseburia* abundance and positively correlated with *Escherichia* abundance under the nominal FDR of 0.25 (Morgan et al., 2015). In this analysis, we took the expression values of complement cascade genes (CFI, C2, and CFB) as the outcomes of interest, and applied the proposed CKF and CKF+ method to detect host gene expression-associated genera for each outcome under the FDR threshold of 0.25. For the initial screening step, we fixed the screening sample size *n*_0_ = 100 with screening set size |*Ŝ*_0_| = 40 as done in simulations. As the data-splitting is random, we repeated the CKF algorithm 10 times with different splits and report those taxa that appeared in more than one of the splits. By using multiple split matrices, we were more likely able to identify any possible signals under the desired FDR level.

In Table 3, we report taxa that were identified as host gene expression associated in our analysis. Taxa in bold were also identified in the original paper (Morgan et al., 2015) using marginal method to control the FDR at 0.25. For the coefficient column of Table 3, we fit the reduced linear regression models with predictors of both selected taxa and clinical variables including disease subtype, antibiotic use, tissue location and inflammatory score, as done previously (Morgan et al., 2015). These clinical variables were included in the model to adjust for potential confounding effects and to obtain a more accurate estimate of the microbiome effect on host gene expression. The sign of a taxon coefficient reflects the direction of association (activation or inhibition). Recall that five taxa *Sutterella, Akkermansia, Roseburia, Bifidobacterium* and *Escherichia* were detected in the original principal component-based marginal analysis (Morgan et al., 2015). All these five except *Roseburia* were identified in our analysis in more than one split. Moreover, we further see that the coefficient signs for each taxa of interest are consistent with the expected direction posited by Morgan et al. (2015). In other words, we correctly identify a majority of taxa of interest function as inhibitors (negative coefficient) or activators (positive coefficient) for each cascade gene expression.

**Table 3.**
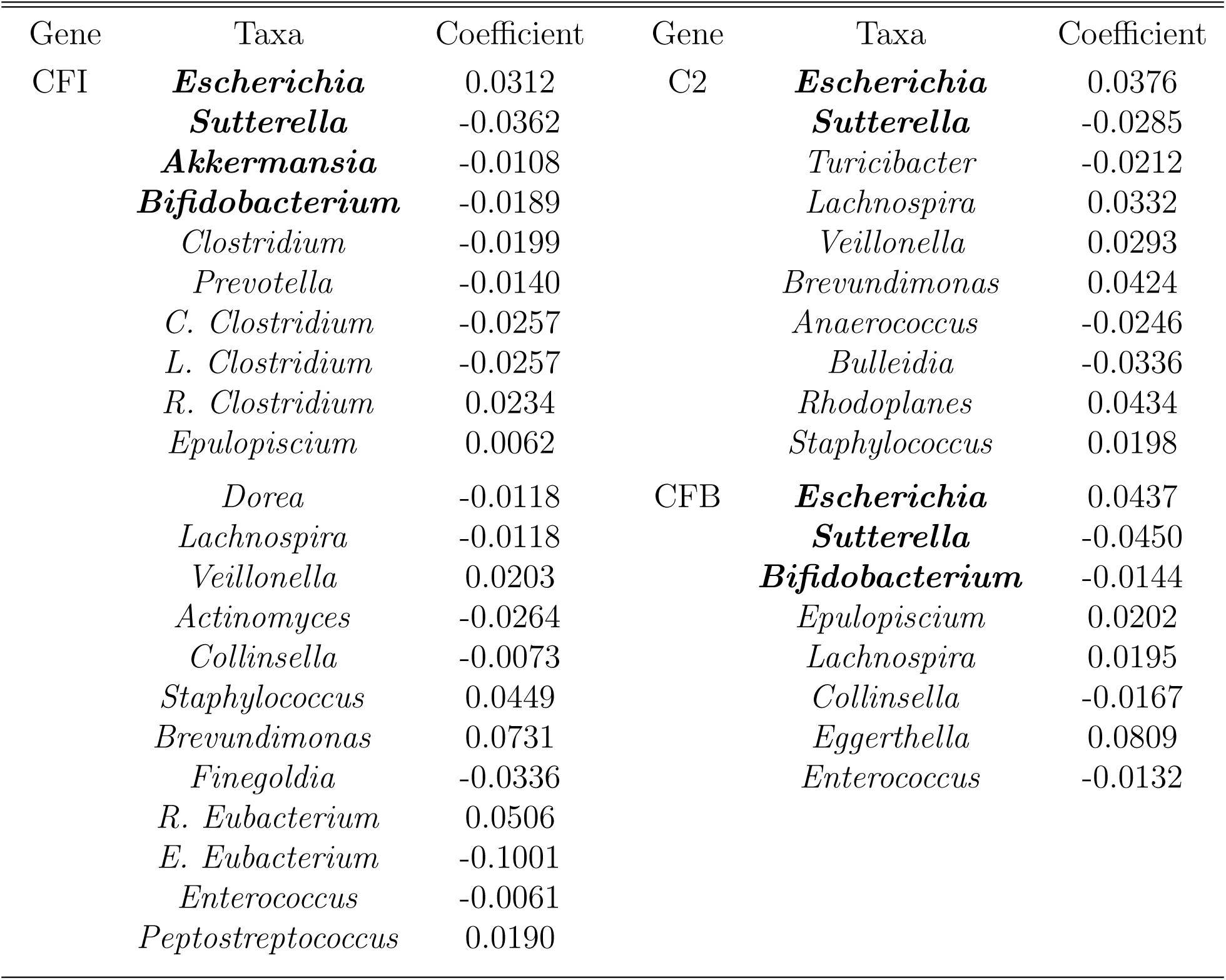
Taxa identified as host gene expression associated under the nominal FDR of 0.25.

We also observe that the taxa set identified for each cascade gene are different, which suggests that specific taxa play key roles on individual gene expression. *Escherichia* and *Sutterella* appear in all gene sets, and *Escherichia* in particular was noted by Morgan et al. (2015) to be hugely influential in patients with inflammatory bowel issues. Despite that we missed taxa *Roseburia* compared to the original analysis, many new taxa were identified as complement cascade gene expression-associated in our CKF analysis. For example, *Epulopiscium* appears in the selection sets for both the CFB and CFI as an inhibitor which may be of particular interest. Likewise, *Lactobacillus* appears in both the CFB gene and C2 gene acting as an activator. On the other hand, *Lachnospira* appears to be an activator for both C2 and CFB but an inihibitor for CFI. The mechanism of how these new taxa affect the host transcript pattern warrants further laboratory investigation.

To conclude, the proposed CKF is more powerful in detecting significant taxa than the original principal component-based marginal analysis (Morgan et al., 2015) under the same nominal FDR of 0.25. Our new method not only provides additional statistical support to results obtained from the original analysis but also gains new biological and biomedical insights on how taxa interact with host complement cascade gene expressions.

## 6. Discussion

In this paper, we consider the problem of identifying outcome-associated microbiome features under a pre-specified FDR. Traditional methods usually cast this problem into a multiple testing framework and examines each microbiome feature individually followed by certain multiple testing procedures to control the FDR. To avoid the potential heavy multiple adjustment burden, we alternatively adopt a joint approach which regresses the response on all microbiome features and achieve FDR control via applying the compositional knockoff filter to the regression. As shown in the numerical studies, the proposed CKF method is more powerful than the marginal BH procedure, and can achieve false discovery control compared to the compositional lasso method. Further numerical study demonstrates a gain in power through employing CKF over the original model-X knockoffs under the logistic normal setting and denser signal scenarios under the Dirichlet-multnomial settings. CKF is extremely useful for microbiome compositional data analysis, as it may be more natural to place the burden of knowledge on the response instead of the features as we have yet been able to develop means to efficiently construct model-X knockoff features for common distributions used for microbiome analysis. Finally, the application our method to the host-microbiome data not only identifies most of gene expression-associated taxa that were identified in the original study (Morgan et al., 2015), but also leads to new discoveries, which may provide new biological insights with further laboratory investigation.

As noted by a referee, a wide array of penalized methods have been proposed for the analysis of high-dimensional regression problems. Methods such as the debiased Lasso (Zhang and Zhang, 2014; van de Geer et al., 2014; Javanmard and Montanari, 2018; van de Geer, 2019) and the MOCE (Wang et al., 2019) method are not guaranteed to retain the compostional constraint on ***β*** under the log-contrast model after the debiasing step. However, it is of future interest to study debiasing methods that retain the sum constraint. The CKF procedure is in the class of “screen and clean” methods such as Wasserman and Roeder (2009) and Meinshausen et al. (2009). However, Meinshausen et al. (2009) does not account for the underlying sparsity assumption in high-dimensional microbiome compositional analysis. Further, these methods do not employ recycling which can lead to a reduction of power, which is especially pronounced in Wasserman and Roeder (2009) which relies on a three-way sample split. Finally, the aforementioned methods do not ensure finite sample FDR control which is a key benefit of the CKF procedure.

Currently, our method can only identify microbial taxa that are associated with a single continuous outcome variable. It is of future interest to extend CKF to more complicated models such as survival models (Plantinga et al., 2017), multivariate-outcome models (Zhan et al., 2017a,b) and generalized linear models (Lu et al., 2019) to accommodate microbiome association studies with more complicated designs. The canonical approach of microbiome fine-mapping is to plug in marginal p-values into the BH procedure to identify outcome-associated taxa under FDR control (Paulson et al., 2013; Parks et al., 2014; Wang and Jia, 2016). Under this vein, there has been a wealth of research interest to utilize additional specific information (e.g., phylogenetic information) of microbiome data to increase the power of detection and maintain control of the FDR (Xiao et al., 2017; Jiang et al., 2017; Hu et al., 2018). It is of future interest to incorporate such information to our CKF framework to further boost the detection power while controlling the FDR at a certain threshold.

## Acknowledgements

The authors wish to thank the editor, associate editor and three referees for their insightful comments and suggestions that have improved the paper. This research was partially supported by the National Institutes of Health grants R21AI144765 and T32GM102057, and National Science Foundation grants DMS-1811552 and DMS-1953189.

## 7. Supporting Information

Supporting Information referenced in Section 3 and Section 4 are available with this article at the *Biometrics* website on Wiley Online Library. It includes discussion on the normalization procedure after CSP, assumptions of Theorem 1 in the context of microbiome data, proofs of lemmas and theorems, additional numerical evaluations, and R code to implement the proposed methods.

## Notes

### Competing Interest Statement

The authors have declared no competing interest.

### Summary of Updates

Main manuscript and supplemental revision

