## Supplemental Materials for "Compositional knockoff filter for high-dimensional regression analysis of microbiome data"

BIOMETRICS 000, 1–30 DOI: 000
000 0000

### Compositional knockoff filter for high-dimensional regression analysis of microbiome data

**Arun Srinivasan[1], Lingzhou Xue[1,*], and Xiang Zhan[2,**]**

[1]Department of Statistics, Pennsylvania State University, University Park, PA 16802, U.S.A.

[2]Department of Public Health Sciences, Pennsylvania State University, Hershey, PA 17033, U.S.A.

SUMMARY: A critical task in microbiome data analysis is to explore the association between a scalar response of interest and a large number of microbial taxa that are summarized as compositional data at different taxonomic levels. Motivated by fine-mapping of the microbiome, we propose a two-step compositional knockoff filter (CKF) to provide the effective finite-sample false discovery rate (FDR) control in high-dimensional linear log-contrast regression analysis of microbiome compositional data. In the first step, we employ the compositional screening procedure to remove insignificant microbial taxa while retaining the essential sum-to-zero constraint. In the second step, we extend the knockoff filter to identify the significant microbial taxa in the sparse regression model for compositional data. Thereby, a subset of the microbes is selected from the high-dimensional microbial taxa as related to the response using a pre-specified FDR threshold. We study the asymptotic properties of the proposed two-step procedure, including both sure screening and effective false discovery control. We demonstrate the finite-sample properties in simulation studies, which show the gain in the empirical power while controlling the nominal FDR. The potential usefulness of the proposed method is also illustrated with application to an inflammatory bowel disease dataset to identify microbial taxa that influence host gene expressions.

KEY WORDS: Compositional constraint; Compositional screening; FDR control; Knockoff filter; Log-contrast model; Microbiome.

This paper has been submitted for consideration for publication in *Biometrics*

#### 1. Introduction

##### 2.1 *Log-Contrast Model*

We use the log-contrast model (Aitchison and Bacon-shone, 1984) for joint microbiome regression analysis. Let $\mathbf{Y} \in \mathbb{R}^n$ denote the response vector and $\mathbf{X} \in \mathbb{R}^{n \times p}$ denote a matrix of microbiome compositions. By structure of the microbiome compositional components, each row of $\mathbf{X}$ must individually sum to 1. Thus $\mathbf{X}$ is not of full rank, leading to identifiability issues for the regression parameters. In order to account for this structure, the log-linear contrast model is often used for compositional data (Aitchison, 2003; Lin et al., 2014). We assume that $X_{ij} > 0$ by replacing the zero proportions by a tiny pseudo positive value as routinely performed in practice (Lin et al., 2014; Shi et al., 2016; Cao et al., 2017; Lu et al., 2019; Zhang et al., 2019). Let $\mathbf{Z}^p \in \mathbb{R}^{n \times (p-1)}$ be the log-ratio transformation of $\mathbf{X}$, where $Z_{ij}^p = \log(X_{ij}/X_{ip})$ and $p$ denotes the reference covariate. The linear log-contrast model is formulated as $\mathbf{Y} = \mathbf{Z}^p \boldsymbol{\beta}_{\setminus p} + \varepsilon$, where $\boldsymbol{\beta}_{\setminus p}$ is the vector of $(p-1)$ coefficients $(\beta_1, \beta_2, ..., \beta_{p-1})$ and error $\varepsilon \sim \mathcal{N}(0, \sigma^2 \mathbf{I})$. To avoid picking a reference component for better model interpretability, the log-contrast model is often reformulated into a symmetric form with a sum-to-zero constraint (Lin et al., 2014). That is,

$$y_i = \sum_{j=1}^{p} Z_{ij}\beta_j + \varepsilon_i \quad \text{subject to} \quad \sum_{j=1}^{p} \beta_j = 0, \tag{1}$$

where $\mathbf{Z} \equiv \{Z_{ij}\}$ is the $n \times p$ log-composition matrix with $Z_{ij} = \log(X_{ij})$ and $\boldsymbol{\beta} \equiv (\beta_1, \beta_2, ..., \beta_p)'$ are the regression coefficients for microbiome covariates. For ease of presentation, model (1) does not explicitly include other covariates, but all the results in the rest of this article still hold in presence of other covariates.

#### 2.2 *Compositional Screening Procedure*

As the fixed-X knockoff requires that $n \geqslant 2p$, screening the predictor set to a low-dimensional setting is necessary for the analysis of high-dimensional compositional data. Let $n_0$ denote the number of samples to use for screening and $n_1$ denote the remaining observations, where $n = n_0 + n_1$. We randomly split the original data $(\mathbf{Z}, \mathbf{Y})$ into $(\mathbf{Z}^{(0)}, \mathbf{Y}^{(0)})$ and $(\mathbf{Z}^{(1)}, \mathbf{Y}^{(1)})$, where $\mathbf{Z}^{(0)} \in \mathbb{R}^{n_0 \times p}$, $\mathbf{Y}^{(0)} \in \mathbb{R}^{n_0}$, $\mathbf{Z}^{(1)} \in \mathbb{R}^{n_1 \times p}$ and $\mathbf{Y}^{(1)} \in \mathbb{R}^{n_1}$. By ensuring that $\mathbf{Z}^{(0)}$ and $\mathbf{Z}^{(1)}$ are disjoint, we are able to implement a recycling step to reuse the original screening data $\mathbf{Z}^{(0)}$, in order to increase the selection power. To this end, we first use the sub-data $(\mathbf{Z}^{(0)}, \mathbf{Y}^{(0)})$ to perform the screening and obtain a subset of features $\hat{S}_0 \subset \{1, ..., p\}$ such that $|\hat{S}_0| \leqslant \frac{n_1}{2}$, where $|\hat{S}_0|$ denotes the cardinality of set $\hat{S}_0$. Throughout this paper, we always assume $|\hat{S}_0| \leqslant \frac{n_1}{2}$ to ensure that we are able to construct the fixed-X knockoffs (Barber and Candès, 2015) for data $(\mathbf{Z}^{(1)}, \mathbf{Y}^{(1)})$ in the subsequent selection step. As the selection step further reduces the feature set after screening, we must ensure that true signals are not lost before the selection step. For this reason, we desire screening methods that attain the sure screening property (Fan and Lv, 2008). That is, with high probability, we desire the selection set estimated by the screening method of choice to contain all relevant features. It is popular to perform screening using Pearson correlation (Fan and Lv, 2008; Fan and Song, 2010; Xue and Zou, 2011) or distance correlation (Li, Zhong and Zhu, 2012). Despite that both marginal correlations-based screening methods enjoy the sure screening property asymptotically, these methods do not account for the compositional nature of microbiome data, which might lead to inefficient inference. We will further demonstrate this issue in the simulation studies of Section 4.1.

To account for the compositional structure, we introduce the novel compositional screening procedure to improve the efficiency for screening microbiome compositional covariates. In general, best-subset selection is often used to identify the optimal $k$ best features (Beale,

Kendall and Mann, 1967). In our log-contrast model, the best-subset selection problem can be expressed as a constrained sparse least-squares estimation problem as follows:

$$\min_{\beta} \frac{1}{2n} ||\mathbf{Y} - \mathbf{Z}\boldsymbol{\beta}||_2^2 \quad \text{s.t.} \quad ||\boldsymbol{\beta}||_0 \leqslant k \quad \text{and} \quad \sum_{j=1}^{p} \beta_j = 0. \tag{2}$$

The proposed compositional screening problem (2) can also be viewed as maximizing the log-likelihood $\ell_n(\boldsymbol{\beta})$ under the sparsity constraint $||\boldsymbol{\beta}||_0 \leqslant k$ (Xu and Chen, 2014). The choice of $k$ is a fundamental question in many high-dimensional screening procedures. Practical domain knowledge may provide information on how sparse one believes the underlying signal is. Common choices for screening set size are often $k = c\lfloor \frac{n_0}{\log(n_0)} \rfloor$ for some $c > 0$ (Fan and Lv, 2008; Li et al., 2012). However, as noted by Li et al. (2012), the choice of screening set size can be viewed as a tuning parameter within the model and concrete means to determine the screening set size are an area of future development. Although (2) is a NP-hard problem, the mixed integer optimization (MIO) allows us to approximately solve the global solution of the nonconvex optimization problem (2) in an efficient manner (Konno and Yamamoto, 2009; Bertsimas, King and Mazumder, 2016). Finally, we demonstrate in the Section 3 that the computed solution of (2) by MIO attains the desirable sure screening guarantees.

After screening, the model reduces to $y_i = \sum_{j \in \hat{S}_0} Z_{ij}\beta_j^r + \varepsilon_i$ subject to $\sum_{j \in \hat{S}_0} \beta_j^r = 0$. Comparing it to the original log-contrast model (1), the regression coefficients in the reduced model $\beta_j^r$ does not necessarily match $\beta_j$ in the original model. To solve this discrepancy, we propose a normalization procedure $X_{ij}^* = X_{ij} / \sum_{j \in \hat{S}_0} X_{ij}$ for $j \in \hat{S}_0$ and for an abuse of notation, we still use $Z_{ij} = log(X_{ij}^*)$ to denote the design matrix to be used in the subsequent selection step. Details about this normalization is available at Section S.1 of the online supporting information.

##### 2.3 *Controlled Variable Selection*

Let $\mathbf{Z}_{\hat{S}_0}^{(1)} \in \mathbb{R}^{n_1 \times |\hat{S}_0|}$ denote the columns of $\mathbf{Z}^{(1)}$ corresponding to $\hat{S}_0$, the selected set from the *computed* solution of (2), and we delineate this from the selection set from the *global* solution of (2) which we instead denote as $\tilde{S}_0$. The knockoff matrix $\tilde{\mathbf{Z}}^{(1)}_{\hat{S}_0}$ is constructed using $\mathbf{Z}^{(1)}_{\hat{S}_0}$ following the fixed-X knockoff framework (Barber and Candès, 2015). The primary assumption of the fixed-X framework is that the burden of knowledge is placed on the design and $\mathbf{Z}^{(1)}$ is assumed to be fixed and the response is generated via a linear Gaussian model. Notably, the fixed-X knockoff places no assumptions on knowing the noise level. Thus, a key appeal of the knockoff filter is the relative lack of strong assumptions needed for theoretical finite-sample control to hold. We refer to Barber and Candès (2015) for a review of the construction of knockoff matrix and for a deeper study into the assumptions needed by the knockoff filter. The use of the screening step allows us to apply the fixed-X knockoff framework in the high-dimensional setting. While fixed-X knockoffs traditionally require a low-dimensional regime, the screening step first reduces the effective dimension to one of size at most $\frac{n_1}{2}$. Thus $\mathbf{Z}^{(1)}_{\hat{S}_0}$ is of dimension at most $n_1 \times \frac{n_1}{2}$. As the knockoff matrix is constructed on $\mathbf{Z}^{(1)}_{\hat{S}_0}$ alone, this satisfies the dimensionality requirements for the construction of the fixed-X knockoff matrix $\tilde{\mathbf{Z}}^{(1)}_{\hat{S}_0}$. To further boost the power of the procedure, we followed data recycling mechanism outlined in Barber and Candès (2019), we construct the recycled knockoff matrix as

$$\tilde{\mathbf{Z}}_{\hat{S}_0} = \begin{bmatrix} \mathbf{Z}^{(0)}_{\hat{S}_0} \\ \tilde{\mathbf{Z}}^{(1)}_{\hat{S}_0} \end{bmatrix}.$$

Note that we treat $(\mathbf{Z}^{(0)}, \mathbf{Y}^{(0)})$ as fixed after the screening step and the first part of knockoff copies are exact copies under the recycling scheme (Barber and Candès, 2019).

We now run the knockoff regression procedure using $\mathbf{Z}_{\hat{S}_0}$, $\tilde{\mathbf{Z}}_{\hat{S}_0}$, and $\mathbf{Y}$. In particular, we first append the screened original and knockoff matrices to create an augmented design matrix $\mathbb{Z}_{\hat{S}_0} = [\mathbf{Z}_{\hat{S}_0}, \tilde{\mathbf{Z}}_{\hat{S}_0}]$. This augmented design matrix is of dimension $\mathbb{Z}_{\hat{S}_0} \in \mathbb{R}^{n \times 2|\hat{S}_0|}$ where the first $|\hat{S}_0|$ features are the original covariates and the remaining $|\hat{S}_0|$ features are the knockoff copies. With this new augmented design matrix, we solve the following Lasso problem:

$$\bar{\boldsymbol{\beta}} = \underset{\boldsymbol{\beta}}{\text{argmin}} \left\{ \frac{1}{2n} ||\mathbf{Y} - \mathbb{Z}_{\hat{S}_0}\boldsymbol{\beta}||_2^2 + \lambda||\boldsymbol{\beta}||_1 \right\} \tag{3}$$

where $\bar{\boldsymbol{\beta}} = (\hat{\boldsymbol{\beta}}, \tilde{\boldsymbol{\beta}})$ is a vector appending the coefficients of original features and knockoff features. Other penalties such as the folded concave penalties (Fan and Li, 2001; Fan et al., 2014) may also be used for the purpose of variable selection. For ease of presentation, we focus on the Lasso problem (3), as existing methods (Lin et al., 2014) do not provide a rigorous FDR control on the selected variables. Comparing to previous problems (1) and (2), we no longer require a sum-to-zero constraint in our augmented Lasso problem (3). This is because, by adding $|\hat{S}_0|$ knockoff features in the augmented design matrix $\mathbb{Z}_{\hat{S}_0}$, the corresponding microbiome data matrix $\mathbb{X}_{\hat{S}_0} = \exp(\mathbb{Z}_{\hat{S}_0})$ is no longer compositional in nature.

The above optimization problem (3) is performed over the entire Lasso path and provides a set of Lasso coefficients denoted by $\{\bar{\boldsymbol{\beta}}(\lambda)\} = \{(\hat{\boldsymbol{\beta}}(\lambda), \tilde{\boldsymbol{\beta}}(\lambda))\}$. Based on $\{\bar{\boldsymbol{\beta}}(\lambda)\}$, we next calculate the knockoff statistic $W_j$, which measures evidence against the null hypothesis $\beta_j = 0$ for each $j \in \hat{S}_0$. For the scope of this paper we use the Lasso signed lambda max statistic (LSM). Let $\mathbf{Z}_{\hat{S}_0,j}$ denote original covariate $j$ and $\tilde{\mathbf{Z}}_{\hat{S}_0,j}$ denote knockoff covariate $j$:

$$W_j(\lambda) = (\max \lambda \text{ such that } \mathbf{Z}_{\hat{S}_0,j} \text{ or } \tilde{\mathbf{Z}}_{\hat{S}_0,j} \text{ enter lasso path}) \times \begin{cases} 1 \text{ if } \mathbf{Z}_{\hat{S}_0,j} \text{ enters before } \tilde{\mathbf{Z}}_{\hat{S}_0,j} \\ -1 \text{ if } \tilde{\mathbf{Z}}_{\hat{S}_0,j} \text{ enters before } \mathbf{Z}_{\hat{S}_0,j} \end{cases} \tag{4}$$

A large and positive $W_j$ would suggest strong evidence that the original feature is significantly outcome-associated as an important feature tends to remain longer in lasso path as $\lambda$ increases. Similarly, a negative or zero $W_j$ value would indicate that the covariate tends to be noise. Thus, $W_j$ is used to calculate the data-dependent knockoff thresholds that ensure finite sample FDR-controlled variable selection. The choice of an $\ell_1$-penalization problem (3) used in the selection step is driven by the need to accurately compute a solution path for construction of the knockoff statistic. As a comparison, the solution of $\ell_0$-penalization problem (2) via MIO does not yield a solution path and is unsuitable for knockoff test statistic construction. Finally, both the standard knockoff and knockoff+ thresholds are considered for the purpose of selection:

Knockoff Threshold:

$$T = \min\left\{t \in \mathcal{W} : \frac{|\{j : W_j \leqslant -t\}|}{1 \vee |\{j : W_j \geqslant t\}|} \leqslant q\right\}, \tag{5}$$

Knockoff+ Threshold:

$$T = \min\left\{t \in \mathcal{W} : \frac{1 + |\{j : W_j \leqslant -t\}|}{1 \vee |\{j : W_j \geqslant t\}|} \leqslant q\right\}, \tag{6}$$

where $q \in [0, 1]$ is the user-specified nominal FDR level, $\mathcal{W} = \{|W_j| : j \in \hat{S}_0\}\backslash\{0\}$ are the unique non-zero values of $|W_j|$'s ($T = +\infty$ if $\mathcal{W}$ is empty) and $a \vee b$ denotes the maximum of $a$ and $b$. Once this threshold has been calculated, we select covariates $\hat{S} = \{j : W_j \geqslant T\}$. Depending on the threshold being used, we term this FDR-control variable selection procedure as either compositional knockoff filter (CKF) or compositional knockoff filter+ (CKF+). For completeness, we summarize the proposed CKF procedures in Algorithm 1.

#### 3. Theoretical Properties

In this section, we first present the theoretical properties of the proposed compositional screening procedure and show that the computed solution from solving the constrained sparse maximum likelihood problem (2) via the mixed integer optimization attains the desired sure screening property. We then summarize the theoretical properties of the proposed compositional knockoff filter method. Leveraging the framework of high-dimensional knockoff filter (Barber and Candès, 2019), we verify that CKF/CKF+ attain finite sample FDR control under the compositional constraint. The main results are presented in this main text and details on the proof to establish these theoretical properties is available through Section S.3 of the online supporting information.

**Algorithm 1 Compositional Knockoff Filter (CKF)**

**Input:** Compositional matrix $\mathbf{X}$ (or log-compositional matrix $\mathbf{Z} = \log(\mathbf{X})$), response $\mathbf{Y}$, FDR threshold $q$, screening sample size $n_0$ and screening set size $|\hat{S}_0|$

**Output:** knockoff selection set $\hat{S}$

**Procedure:**

(1) Randomly split the data $(\mathbf{Z}, \mathbf{Y})$ into disjoint $(\mathbf{Z}^{(0)}, \mathbf{Y}^{(0)})$ and $(\mathbf{Z}^{(1)}, \mathbf{Y}^{(1)})$.

(2) **Screening Step:**

(a) Run the compositional screening procedure method on $(\mathbf{Z}^{(0)}, \mathbf{Y}^{(0)})$ to identify $\hat{S}_0$.

(b) Apply the normalization procedure $X^*_{ij} = X_{ij} / \sum_{j \in \hat{S}_0} X_{ij}$ for $j \in \hat{S}_0$ and calculate $Z_{ij} = log(X^*_{ij})$ as the design matrix to be used in the subsequential selection step.

(3) **Selection Step:**

(a) Generate the recycled knockoff matrix $\tilde{\mathbf{Z}}_{\hat{S}_0}$ and construct the augmented design matrix: $\mathbb{Z}_{\hat{S}_0} = [\mathbf{Z}_{\hat{S}_0} \ \tilde{\mathbf{Z}}_{\hat{S}_0}]$.

(b) Solve equation (3) to calculate the coefficients $\bar{\boldsymbol{\beta}}(\lambda)$.

(c) Calculate knockoff statistics $W_j$ from $\bar{\beta}_j(\lambda)$.

(d) Use the knockoff or knockoff+ threshold (5) and (6) to calculate $T$ from $\mathcal{W}$.

(e) Determine the knockoff or knockoff+ selection set as $\hat{S} = \{j : W_j \geqslant T\}$.

#### 3.1 *Theoretical Properties of Compositional Screening*

We will show in this section that the compositional screening procedure attains the sure screening property. For ease of presentation, some notation is introduced first. Let $s$ denote an arbitrary subset of $\{1, \ldots, p\}$ corresponding to a sub-model with coefficients $\beta_s$, and $S^*$ be the true model with $p^*$ nonzero coefficients, with corresponding true coefficient vector $\boldsymbol{\beta}^*$. Let $\hat{S}_0$ denote the computed screened sub-model after applying the compositional screening procedure. Assume that $\hat{S}_0$ retains at most $k$ features with $p^* < k < p$. Let $\mathbf{S}^k_+ = \{s : S^* \subset$

$s; ||s||_0 \leqslant k\}$ denote the set of all overfit models and $\mathbf{S}_-^k = \{s : S^* \not\subset s; ||s||_0 \leqslant k\}$ denote the set of underfit models. We will show that the compositional screening procedure does not miss true signals with high probability. That is:

$$P(S^* \subset \hat{S}_0) \to 1 \text{ as } n \to \infty. \tag{7}$$

For the technical aspects of our sure-screening proof to hold, we make the following assumptions (1-4), encompassing requirements on the dimension, signal strength and microbiome design matrix:

ASSUMPTION 1: $\log(p) = O(n^m)$ for some $0 \leqslant m < 1$.

ASSUMPTION 2: There exists $w_1 > 0$ and $w_2 > 0$ and non-negative constants $\tau_1$ and $\tau_2$ such that $\min\limits_{j \in S^*} |\beta_j^*| \geqslant w_1 n^{-\tau_1}$ and $p^* < k \leqslant w_2 n^{\tau_2}$.

ASSUMPTION 3: There exist constants $c_1 > 0$ and $\delta_1 > 0$ such that for sufficiently large $n$ such that $\lambda_{min}[n^{-1} \sum_{i=1}^n \mathbf{Z}_{is}\mathbf{Z}_{is}^t] \geqslant c_1$ for $s \in \mathbf{S}_+^{2k}$ and $||\boldsymbol{\beta}_s - \boldsymbol{\beta}_s^*||_2 \leqslant \delta_1$, where $\lambda_{min}[M]$ denotes the smallest eigenvalue of the matrix $M$, and $\mathbf{Z}_{is} = (Z_{ij})_{j \in s}$.

ASSUMPTION 4: There exist constants $c_2 > 0$ and $c_3 > 0$ such that $|Z_{ij}| \leqslant c_2$ and $\max\limits_{1 \leqslant j \leqslant p} \max\limits_{1 \leqslant i \leqslant n} \{\frac{Z_{ij}^2}{\sum_{i=1}^n Z_{ij}^2 \sigma_i^2}\} \leqslant c_3 n^{-1}$ when $n$ is sufficiently large, where $\sigma_i^2 = Var(\mathbf{Y}|\mathbf{Z})$.

Assumption 1 places a weak restriction on $p$ and $n$ of the data, which is very likely to be met in many microbiome studies (Wang and Jia, 2016). Assumption 2 places a restraint on the minimum strength of true signals, such that they are discoverable. This assumption is common for statistical screening and variable selection methods (Fan and Lv, 2008; Fan and Song, 2010; Lin et al., 2014). Assumption 3 corresponds to the UUP condition (Candes and Tao, 2007) which controls the pairwise correlations between the columns of $\mathbf{Z}$. This condition is prevalent across many high-dimensional variable selection methods such as the Dantzig selector (Candes and Tao, 2007), SIS-DS (Fan and Lv, 2008), forward regression (Wang,

2009), and the sparse MLE (Xu and Chen, 2014). We have conducted numerical studies to evaluate the applicability of Assumption 3 in the context of microbiome data settings in Section S.2 of the online supporting information. Finally, as noted by Xu and Chen (2014) and Chen and Chen (2012), Assumption 4 likely will hold for a wide class of design matrices as long $\sigma_i^2$ is not degenerate. In Section S.2 of the online supporting information, we illustrate the validity of Assumption 4 on the mucosal microbiome data analyzed later in this paper. Further details on these assumptions are available in Section S.2 of the online supporting information. Under Assumptions 1–4, Theorem 1 shows that the proposed compositional screening procedure attains the sure screening property. The proof of Theorem 1 relies on two key lemmas which are presented first.

Lemma 1: *Let $\tilde{S}_0$ denote the index set of screened features from the global solution of the constrained sparse maximum-likelihood estimation problem* (2)*, where $|\tilde{S}_0| = k$. Let $\mathbf{S}_+^k = \{s : S^* \subset s; ||s||_0 \leqslant k\}$. Assume that Assumptions 1-4 hold and $\tau_1 + \tau_2 < \frac{(1-m)}{2}$. Then:*

$$P(\tilde{S}_0 \in \mathbf{S}_+^k) \to 1 \text{ as } n \to \infty$$

Lemma 1 ensures that the model selected by the solution of the constrained sparse maximum-likelihood estimation will be in the set of overfit models with high-probability. Thus, this ensures no signals are lost during screening. In other words, the global solution of the constrained sparse maximum-likelihood estimation problem attains the sure screening property.

Lemma 2: *Let $\hat{\boldsymbol{\beta}}_{MIO}$ denote the computed coefficient magnitudes of the model selected by the compositional screening procedure through mixed integer optimization and $\tilde{\boldsymbol{\beta}}$ denote the coefficients of the global solution of the constrained sparse maximum likelihood problem. Given $\varepsilon > 0$, then:*

$$P(||\hat{\boldsymbol{\beta}}_{MIO} - \tilde{\boldsymbol{\beta}}||_\infty < \varepsilon) \to 1$$

Lemma 2 demonstrates that the computed solution of the compositional screening procedure through mixed integer optimization converges to the global solution of the constrained sparse maximum likelihood problem with high probability. By combining Lemma 1 and Lemma 2, it follows that the computed solution attains the sure screening property. This result is presented in Theorem 1.

THEOREM 1: *Given we have $n$ independent observations with $p$ possible features. Assume that Assumptions 1-4 hold and $\tau_1 + \tau_2 < \frac{(1-m)}{2}$. Let $\hat{S}_0$ denote the computed screened set from the compositional screening procedure where $p^\star < |\hat{S}_0| < p$. Then:*

$$P(S^* \subset \hat{S}_0) \to 1 \text{ as } n \to \infty$$

Theorem 1 allows us to claim that the compositional screening procedure will not lose any signals during screening with high probability. In summary, the compositional screening procedure accounts for the compositional constraint and also ensures the screening power.

##### 3.2 *FDR Control Properties of Compositional Knockoff Filter*

In this section, we briefly outline the FDR control properties of CKF. In order to control FDR, the knockoff statistic must obey the *anti-symmetry* and *sufficiency* properties while the design matrix and response must satisfy both the *Pairwise Exchangeability for the Response Lemma* and *Pairwise Exchangeability for the Features Lemma* (Barber and Candès, 2015). In this paper we primarily focus on the LSM knockoff statistic (4) which has been shown to satisfy the *anti-symmetry* and *sufficiency* properties (Barber and Candès, 2019). Therefore, the FDR control properties of CKF are a direct consequence of the FDR control theory outlined in Barber and Candès (2015) and Barber and Candès (2019) and we reiterate the FDR control property here for posterity. As we have validated that the CSP attains the sure-screening property, the compositional knockoff+ threshold ensures finite sample FDR control as stated in the following Theorem 2.

Theorem 2: *For $q \in [0,1]$, the knockoff+ method with data-recycling ensures:*

$$\mathbb{E}\left[\frac{|\{j : \beta_j = 0 \text{ and } j \in \hat{S}\}|}{|\hat{S}| \vee 1}\,\middle|\, E\right] \leqslant q$$

*where $S$ denotes the index set of selected coefficients through the compositional knockoff+ procedure, $E$ denotes the event $\{S^* \subset \hat{S}_0\}$. The expectation is over the Gaussian noise vector $\varepsilon$ and $\mathbf{Z}$ and $\tilde{\mathbf{Z}}$ are fixed.*

Theorem 2 demonstrates that CKF+ controls the FDR at a user-specified level $q$, after conditioning on the results of the screening procedure. By the argument in Theorem 2 of Barber and Candès (2019), if a proper screening procedure which attains the sure screening property (such as our proposed compositional screening procedure through mixed integer optimization) is implemented in the screening step, FDR is controlled even without conditioning on $E$.

Remark 1: Given the above exchangeability results and the previous theorems, the standard knockoff threshold controls a modified form of false discovery rate (Barber and Candès, 2015). In particular, for $q \in [0,1]$, the knockoff method ensures:

$$\mathbb{E}\left[\frac{|\{j : \beta_j = 0 \text{ and } j \in \hat{S}\}|}{|\hat{S}| + q^{-1}}\,\middle|\, E\right] \leqslant q.$$

Compared with the formula in Theorem 2, the additional $q^{-1}$ in the denominator sometimes favors a larger selected set $\hat{S}$ in CKF compared to CKF+. But when the selected set $\hat{S}$ is relatively large or when the nominal FDR threshold $q$ is relatively large, the difference between CKF and CKF+ vanishes as $q^{-1}$ has little effect compared to $|\hat{S}|$ under such scenarios.

Next, we varied the sparsity levels $|S^*| \in \{15, 20, 25, 30\}$ and set the first 30 entries $\boldsymbol{\beta}_{1:30}$ of the whole regression coefficient vector $\boldsymbol{\beta}_{1:400}$ as: $\boldsymbol{\beta}_{1:30} = (-3, 3, 2.5, -1, -1.5; 3, 3, -2, -2, -2; 1, -1, 3, -2, -1; -1, 1, 2, -1, -1; 3, 3, -3, -2, -1; 3, 3, -3, -2, -1)$. The remaining regression coefficients $\boldsymbol{\beta}_{31:400}$ were all set to be zeros. We constructed the regression coefficients in the aforementioned way such that $\sum_{j=1}^{|S^*|} \beta_j = 0$, for each $|S^*| \in \{15, 20, 25, 30\}$. Under this scheme, it is easy to check that the coefficient vector always satisfies the sum-to-zero constraint under each of the four sparsity levels. Finally, we simulated the response vector $\mathbf{Y}$ from $\mathbf{Y} = \mathbf{Z}\boldsymbol{\beta}_{S^*} + \varepsilon$, where $\boldsymbol{\beta}_{S^*} = \boldsymbol{\beta}_{1:|S^*|}, |S^*| \in \{15, 20, 25, 30\}$ and $\varepsilon \sim \mathcal{N}(0, I)$.

$$\widehat{\mathrm{FDR}} = \mathbb{E}_N\left[\frac{|\{j : \beta_j = 0 \text{ and } j \in \hat{S}\}|}{|\hat{S}| \vee 1}\right]; \qquad \widehat{\mathrm{Power}} = \mathbb{E}_N\left[\frac{|\{j : \beta_j \neq 0 \text{ and } j \in \hat{S}\}|}{|S^*|}\right],$$

where $\mathbb{E}_N$ denotes the empirical average over $N = 200$ replicates. The results of empirical FDR and empirical power are reported in Table 2.

**Table 1**
*Average screening proportions of true signals based on 200 replicates under the Dirichlet-multnomial (DM) distribution and logistic normal (LN) distribution.*

| Distribution | Screening Method | $\lvert S^*\rvert = 15$ | $\lvert S^*\rvert = 20$ | $\lvert S^*\rvert = 25$ | $\lvert S^*\rvert = 30$ |
|---|---|---|---|---|---|
| DM | CSP | 1.000 | 1.000 | 1.000 | 1.000 |
| | PC | 0.599 | 0.497 | 0.495 | 0.447 |
| | DC | 0.561 | 0.462 | 0.464 | 0.413 |
| LN | CSP | 0.994 | 0.991 | 1.000 | 1.000 |
| | PC | 0.663 | 0.577 | 0.480 | 0.442 |
| | DC | 0.653 | 0.566 | 0.467 | 0.425 |

**Table 2**
*Empirical FDR and power under nominal FDR of 0.1 based on 200 replicates.*

| Distribution | Metric | Method | $\lvert S^*\rvert = 15$ | $\lvert S^*\rvert = 20$ | $\lvert S^*\rvert = 25$ | $\lvert S^*\rvert = 30$ |
|---|---|---|---|---|---|---|
| DM | $\widehat{\text{FDR}}$ | CKF | 0.132 | 0.107 | 0.102 | 0.102 |
| | | CKF+ | 0.073 | 0.064 | 0.070 | 0.075 |
| | | KF | 0.122 | 0.117 | 0.110 | 0.108 |
| | | KF+ | 0.068 | 0.084 | 0.079 | 0.084 |
| | | CL | 0.814 | 0.783 | 0.670 | 0.620 |
| | | BH | 0.106 | 0.095 | 0.100 | 0.102 |
| | $\widehat{\text{Power}}$ | CKF | 0.954 | 0.961 | 0.968 | 0.968 |
| | | CKF+ | 0.881 | 0.907 | 0.946 | 0.935 |
| | | KF | 0.999 | 0.998 | 0.953 | 0.931 |
| | | KF+ | 0.990 | 0.974 | 0.881 | 0.851 |
| | | BH | 0.626 | 0.547 | 0.445 | 0.385 |
| LN | $\widehat{\text{FDR}}$ | CKF | 0.132 | 0.107 | 0.102 | 0.102 |
| | | CKF+ | 0.073 | 0.064 | 0.070 | 0.075 |
| | | KF | 0.101 | 0.115 | 0.101 | 0.090 |
| | | KF+ | 0.064 | 0.070 | 0.062 | 0.054 |
| | | CL | 0.825 | 0.797 | 0.778 | 0.765 |
| | | BH | 0.094 | 0.108 | 0.097 | 0.087 |
| | $\widehat{\text{Power}}$ | CKF | 0.954 | 0.961 | 0.968 | 0.968 |
| | | CKF+ | 0.881 | 0.907 | 0.946 | 0.935 |
| | | KF | 0.849 | 0.755 | 0.691 | 0.577 |
| | | KF+ | 0.730 | 0.582 | 0.555 | 0.457 |
| | | BH | 0.521 | 0.426 | 0.475 | 0.409 |

**Table 3**
*Taxa identified as host gene expression associated under the nominal FDR of 0.25.*

| Gene | Taxa | Coefficient | Gene | Taxa | Coefficient |
|---|---|---|---|---|---|
| CFI | ***Escherichia*** | 0.0312 | C2 | ***Escherichia*** | 0.0376 |
| | ***Sutterella*** | -0.0362 | | ***Sutterella*** | -0.0285 |
| | ***Akkermansia*** | -0.0108 | | *Turicibacter* | -0.0212 |
| | ***Bifidobacterium*** | -0.0189 | | *Lachnospira* | 0.0332 |
| | *Clostridium* | -0.0199 | | *Veillonella* | 0.0293 |
| | *Prevotella* | -0.0140 | | *Brevundimonas* | 0.0424 |
| | *C. Clostridium* | -0.0257 | | *Anaerococcus* | -0.0246 |
| | *L. Clostridium* | -0.0257 | | *Bulleidia* | -0.0336 |
| | *R. Clostridium* | 0.0234 | | *Rhodoplanes* | 0.0434 |
| | *Epulopiscium* | 0.0062 | | *Staphylococcus* | 0.0198 |
| | *Dorea* | -0.0118 | CFB | ***Escherichia*** | 0.0437 |
| | *Lachnospira* | -0.0118 | | ***Sutterella*** | -0.0450 |
| | *Veillonella* | 0.0203 | | ***Bifidobacterium*** | -0.0144 |
| | *Actinomyces* | -0.0264 | | *Epulopiscium* | 0.0202 |
| | *Collinsella* | -0.0073 | | *Lachnospira* | 0.0195 |
| | *Staphylococcus* | 0.0449 | | *Collinsella* | -0.0167 |
| | *Brevundimonas* | 0.0731 | | *Eggerthella* | 0.0809 |
| | *Finegoldia* | -0.0336 | | *Enterococcus* | -0.0132 |
| | *R. Eubacterium* | 0.0506 | | | |
| | *E. Eubacterium* | -0.1001 | | | |
| | *Enterococcus* | -0.0061 | | | |
| | *Peptostreptococcus* | 0.0190 | | | |
